# Reduced inter-subject functional connectivity during movies in autism: Replicability across cross-national fMRI datasets

**DOI:** 10.1101/2025.02.04.636405

**Authors:** Feng Lin, Laura Albantakis, Tuomo Noppari, Severi Santavirta, Marie-Luise Brandi, Lihua Sun, Lasse Lukkarinen, Pekka Tani, Juha Salmi, Lauri Nummenmaa, Juergen Dukart, Leonhard Schilbach, Juha M. Lahnakoski

## Abstract

**Background:** Autism is a neurodevelopmental disorder characterized by repetitive behaviors and difficulties in social communication and interaction. Previous research has shown that these symptoms are linked to idiosyncratic behavioral and brain activity patterns while viewing natural social events in movies. This study aimed to investigate the replicability of brain activity idiosyncrasy in autistic individuals by comparing their inter-subject functional connectivity (ISFC) with that of neurotypical individuals.

**Methods:** We tested for ISFC differences between autistic and neurotypical groups using functional magnetic resonance imaging (fMRI) data from two independent datasets from Germany (N_neurotypical_ = 25, 7 Males, 18 Females; N_autism_ = 22, 12 Males, 10 Females) and Finland (N_neurotypical_ = 19, N_autism_ = 18; All males). Participants watched short movie stimuli, and pairwise ISFCs were computed across 273 brain regions. Group differences were evaluated using subject-wise permutation tests for each dataset.

**Results:** In both datasets, the autistic group showed lower ISFCs compared to the neurotypical group, specifically between visual regions (e.g., occipital gyrus, cuneus) and parietal regions (e.g., superior and inferior parietal lobules), as well as between visual regions and frontal regions (e.g., inferior frontal gyrus, precentral gyrus). ISFC was higher in the Finnish autistic group in temporal regions associated with sound and speech processing.

**Conclusions:** The study confirmed the replicability of reduced ISFCs in autistic individuals during naturalistic movie-watching, especially between visual and parietal/frontal brain regions. These findings reinforce the utility of ISFC and naturalistic movie-watching paradigm in studying neural connectivity alterations in autism.

## 1 Introduction

Autism is a neurodevelopmental disorder characterized by difficulties in social communication, interaction, and repetitive behaviors with restricted interest. Altered sensory responses to external stimuli have been suggested as a potential underlying mechanism (1). Research has shown that autistic individuals exhibit significant variability in symptom severity, particularly in in social impairments and overall functioning (2-4). While many autistic individuals without intellectual impairment perform well in controlled tasks, such as recognizing emotional facial expressions (5-7), they may struggle in more naturalistic or socially dynamic settings (8,9). These findings underscore the need to study individual differences in autism within more ecologically valid scenarios.

Naturalistic functional magnetic resonance imaging (fMRI) provides an effective way to examine ‘social brain’ activity in dynamic, real-world (10,11). In this paradigm, participants are presented with complex audiovisual stimuli, such as movies, without performing a specific task. Studies using naturalistic viewing paradigms have reported reduced neural similarity in autistic compared to neurotypical individuals (2, 12-15). Neural similarity has been associated with participants’ family relations (16), friendships (17), and psychological perspectives (18-20). A common measure of neural similarity in naturalistic fMRI is inter-subject correlation (ISC), which assesses regional brain activity synchronization across individuals during movie-watching (21). Expanding on this, inter-subject functional connectivity (ISFC) compares fMRI time courses between different brain regions across individuals (22). ISFC distinguishes stimulus-dependent interregional correlations from stimulus-independent correlations, i.e. intrinsic brain activity and noise, such as head motion and physiological artifacts. It therefore provides a more refined measure of neural similarity. ISFC has shown promise in detecting functional differences across groups, such as tracking symptom progression in psychiatric conditions like schizophrenia (23).

Previous studies have reported lower ISC and increased occurrence of atypical brain network states in autism compared to neurotypical controls (2, 17-19). Specifically, one study using naturalistic movie fMRI found that young autistic individuals exhibited reduced ISFC, particularly in visual, sensorimotor, and subcortical networks (24). These findings suggest that ISFC patterns may differ between autistic and neurotypical individuals. Given the ongoing “replication crisis” in neuroimaging (25,26), it is crucial to assess the reproducibility of fMRI findings. Additionally, cross-cultural factors have been shown to contribute to the significant inter-subject variability in autistic individuals (27). Inter-subject variability within the autistic group can be indicated by either increased (intersubject hyperconnectivity) or decreased (intersubject hypoconnectivity) ISFC when compared to the neurotypical group. Our study aimed to investigate the inter-subject variability in autism by comparing ISFC between autistic and neurotypical individuals. We also sought to determine whether the results could be replicated across cross-national fMRI datasets, collected both with a similar naturalistic paradigm and analyzed using identical preprocessing and statistical pipelines. Based on previous research, we hypothesized that ISFC in autistic individuals would be lower than in neurotypical individuals, particularly in brain regions associated with social cognition, such as the temporoparietal junction, superior temporal sulcus, precuneus, medial prefrontal cortex, and fusiform gyrus, as suggested by the ‘social brain’ concept (28,29).

## 2 Methods and Materials

### 2.1 Participants

Here, we evaluated independent German and Finnish datasets. The German dataset contained 22 autistic adults (Mean age 35.72 ± 10.58 years, range 20-53 years, 12 males) and 25 neurotypical participants (Mean age 32.76 ± 12.20 years, range 18-60 years, 7 males). Data was collected at Max Planck Institute of Psychiatry, Munich, Germany, and protocols were approved by the ethics committee of the Ludwig-Maximilians-University (LMU) Munich. All procedures were performed in accordance with the declaration of Helsinki. All subjects gave a written informed consent prior to their participation.

The Finnish dataset contained 20 autistic adults (Mean age 27.25 ± 5.72 years, range 20-40, only males) and 19 neurotypical participants (Mean age 28.52 ± 7.69 years, range 20-47, only males). Data were collected at Turku University Hospital, Turku, Finland. The anatomical findings and results of emotional face perception task from this dataset have been published in recent studies (30,31). The study was approved by the ethics committee of the Hospital District of Southwest Finland and was conducted in accordance with the declaration of Helsinki. All subjects provided informed consent prior to the study.

### 2.2 Stimuli and Procedure

In the German dataset, fMRI data were collected while participants watched 52 short (9-22s) clips of Hollywood movies depicting socio-emotional scenes across five categories (i.e., emotion, neutral, social interaction, non-interaction, pain) that were selected from a database of 137 videos described in more detail in a previous study (32). The clips were played without sound to avoid confounds due to different proficiency in English among German participants. In the Finnish dataset, fMRI data were collected using a similar naturalistic movie-watching paradigm, with 54 short (9-22s) clips depicting the same categories as in the German sample. Finnish people generally demonstrate a strong proficiency in English, attributed to a minimum of seven years of compulsory education in the language. Thus, the soundtracks of videos were retained during the Finnish experiment since the participants were assumed to understand dialogues in the movies sufficiently. The total duration of movie clips in both datasets is approximately 11 minutes. The clinical characteristics of all individuals are described and listed in Supplementary Tables 1-4.

### 2.3 Image acquisition and Preprocessing

The German whole-brain structural and fMRI data were acquired on a GE Discovery MR750 3T scanner. Anatomical brain images were collected using a T1-weighted (T1w) sequence (TR = 6.2 ms, TE = 2.3 ms) with 1 mm^3^ isotropic voxel size. The fMRI data were collected with an echo-planar imaging sequence (TR = 2000 ms, TE = 20 ms, flip angle = 90°, 400 mm FOV, 128 × 128 reconstruction matrix, 3.5 mm slice thickness).

The Finnish whole-brain structural and fMRI data were collected using a Phillips Ingenuity TF PET/MR 3T scanner. Structural brain images were acquired using a T1w sequence (TR = 9.8 ms, TE = 4.6 ms, flip angle = 7°, 250 mm FOV, 256 × 256 reconstruction matrix) with 1 mm^3^ isotropic voxel size. Functional data were collected with a T2*-weighted echo-planar imaging sequence (TR = 2600 ms, TE = 30 ms, flip angle = 75°, 240 mm FOV, 80 × 80 reconstruction matrix, 3.0 mm slice thickness).

A whole-brain atlas with 273 regions of interests (ROIs) from Brainnetome (33) combined with a probabilistic atlas of the human cerebellum (34,35) were used to extract regional BOLD time series from the voxel-wise whole brain fMRI. Structural T1w images and fMRI data were preprocessed using fMRIPrep 1.3.0.2 (36). Additional preprocessing steps were performed with Nilearn 0.10.1 (https://github.com/nilearn/nilearn, 37) to control for nuisance variables and low-frequency signal components estimated by fMRIPrep. High-pass filtering was conducted via adding discrete cosine transformation basis regressors from fMRIPrep confound outputs. Additionally, linear trends of signals were removed. Signal artifacts were handled through linear confound removal, using eight parameters: the average signals from white matter and cerebrospinal fluid, along with six basic motion parameters (translation/rotation). In the end, time series were shifted to zero mean and scaled to unit variance, using the sample standard deviation. After checking framewise displacement (FD) and standardized DVARS (38), two subjects in the Finnish autism group were excluded from further analysis due to high average FD values (threshold = 0.5). A total of 47 subjects were analyzed in the German dataset (N_autism_ = 22, N_neurotypical_ = 25) and 37 subjects in the Finnish dataset (N_autism_ = 18, N_neurotypical_ = 19).

### 2.4 Inter-subject functional connectivity

ISFCs were calculated separately for each group (German/Finnish, Autism/Neurotypical) using the Python package BrainIAK (Brain Imaging Analysis Kit, http://brainiak.org). Based on the similar concepts of ISC (correlation between same region across brains) and ISFC (correlation between different regions across brains), customized Python scripts were adapted from the ISC analysis part of the BrainIAK package. Pairwise ISFC calculations were conducted between all pairs of the 273 ROIs. Since we did not analyze the directionality of the connections, we calculated symmetrical connectivity matrices by averaging each matrix with its transpose. After this, a total of 37,128 pairs of unique median ISFC values were calculated from the time series of each group. Subsequently, an ISFC matrix with group differences was created.

### 2.5 Statistical analysis

Non-parametric methods were used for statistical testing of ISFC to account for the non-normal value distribution and ensure robust hypothesis testing without parametric assumptions (39). Median values were computed as summary statistic as suggested for group comparison (40). To compare the difference between autism and neurotypical individuals within each dataset, subject-wise permutation was used for the Finnish and German datasets (neurotypical-autism). During permutations, pairwise median ISFC differences were used as the summary statistics and 5000 iterations with randomized group labels were implemented for the analysis of each dataset with BrainIAK. A pairwise ISFC group difference was considered statistically significant if its magnitude exceeded the 97.5th percentile of the null distribution derived from permutation testing, corresponding to p < .05 (two-sided). For visualization purposes, significant ISFC group differences at more stringent threshold (p < .01, two-sided) were also calculated. Further discussion on the validation of the subject-wise permutation method is provided in the Discussion section. Due to the systematic difference in the sex distribution of participants between the two datasets, we evaluated the replicability of the ISFC group differences between the male and female groups in the German data. Additionally, we compared German males and females to the Finnish dataset, which contained only males. We also evaluated whether there are age and sex effects on ISFC group differences across the two datasets (see Supplemental results).

To calculate the replication rate of ISFC group differences (neurotypical-autism) across two datasets, the percentage of replicated significant pairwise ISFCs from the German and Finnish datasets was counted. These pairwise ISFCs were identified at p < .05 in each dataset for the calculation of replicated ISFC group differences. Specifically, only pairwise ISFCs with identical pairs of ROI indices were considered as “replicated” results. The replication rate was determined by computing the proportion of significant pairwise ISFC differences in the German dataset that also occurred in the Finnish dataset, relative to the total number of significant ISFC pairs in the German dataset. The German dataset was used as the discovery data and the Finnish dataset as the replication data because the German dataset lacked auditory stimulation. Thus, we did not expect all effects observed in the Finnish data to replicate in the German sample. The significance of the replication rate was evaluated by randomly shuffling the ROI order (both rows and columns shuffled in the same order) 5000 times to produce a null distribution. The analysis steps are illustrated in Figure 1.

**Figure 1.**
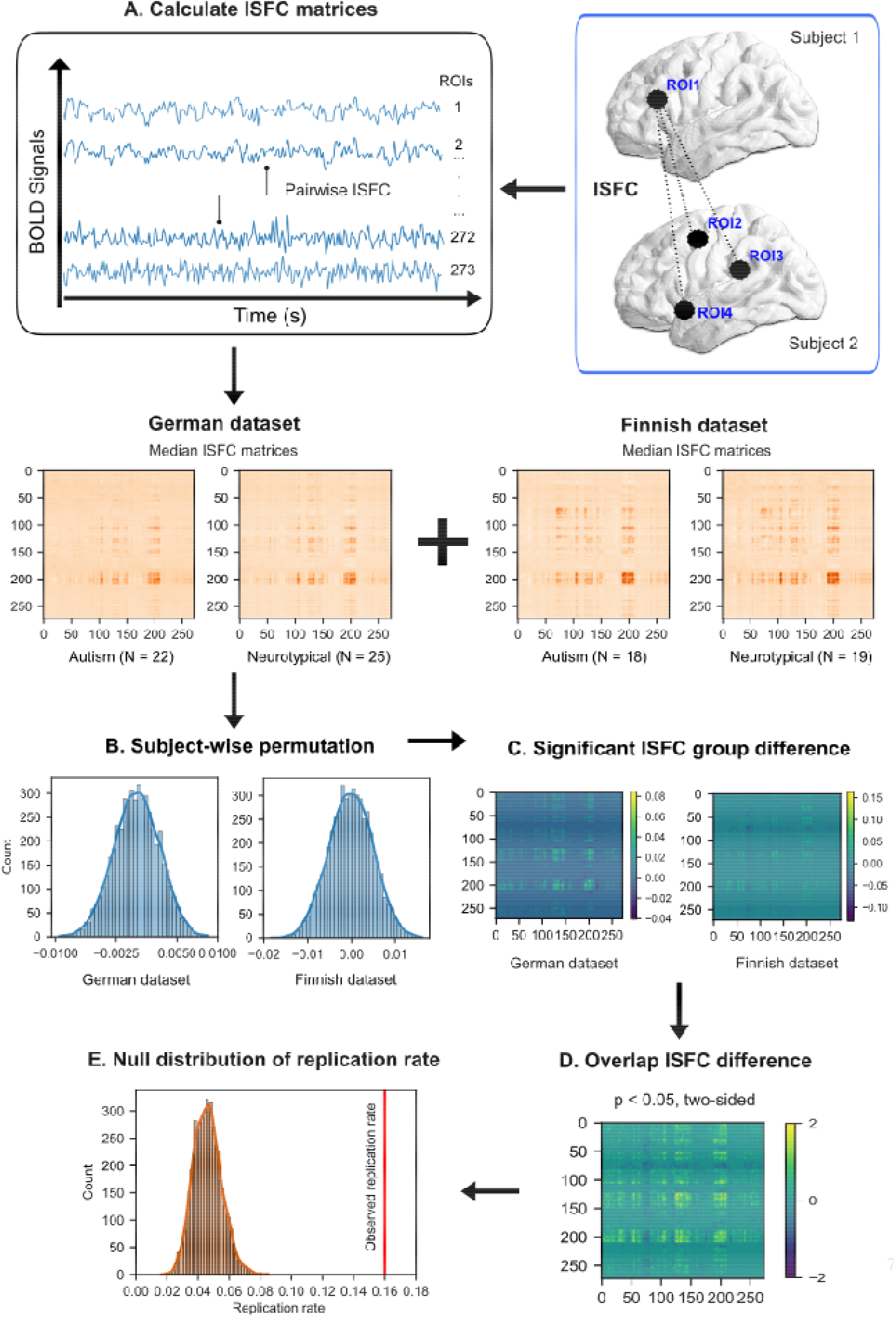
Methods. (A) ISFC matrices were calculated for four participant groups. Median pairwise ISFC values were obtained for each participant group. (B) Subject-wise permutation were conducted 5000 times for the group difference of ISFC in the German and Finnish dataset.(C) The significant ISFC group differences in the two datasets were calculated based on the permutation distributions. (D) The overlapping ISFC difference across two datasets were computed as the intersection of the two matrices from step C. (E) Null distribution of replication rate ISFC across two datasets. The significance of the actual replication rate was compared against repeated analyses of replication rates with permuted ROI orders.

## 3 Results

### 3.1 ISFC distribution across different groups and datasets

The median pairwise ISFCs among the 273 ROIs for the two groups and two datasets are presented in Figure 2A. Overall, the Finnish dataset showed higher maximum ISFCs for both groups compared to the German dataset. In the German data, the maximum ISFC for the neurotypical group (max=0.359) was higher than for the autism group (max=0.329), whereas the opposite was observed in the Finnish data (max=0.462 and 0.474, respectively). The ISFC differences between neurotypical and autism groups were significantly correlated between the two datasets (Pearson r=0.156, p_permuted_ < .001), although the group difference was overall larger in the Finnish dataset than in the German dataset. The highest ISFC difference reached 0.162 in the Finnish dataset, while the maximal difference in the German dataset was 0.084 (Figure 2B). Significant ISFCs were observed within and between occipital and parietal/frontal regions in both groups across both datasets. Additionally, in the Finnish data, significant ISFCs extended to temporal regions (Figure 2C).

**Figure 2.**
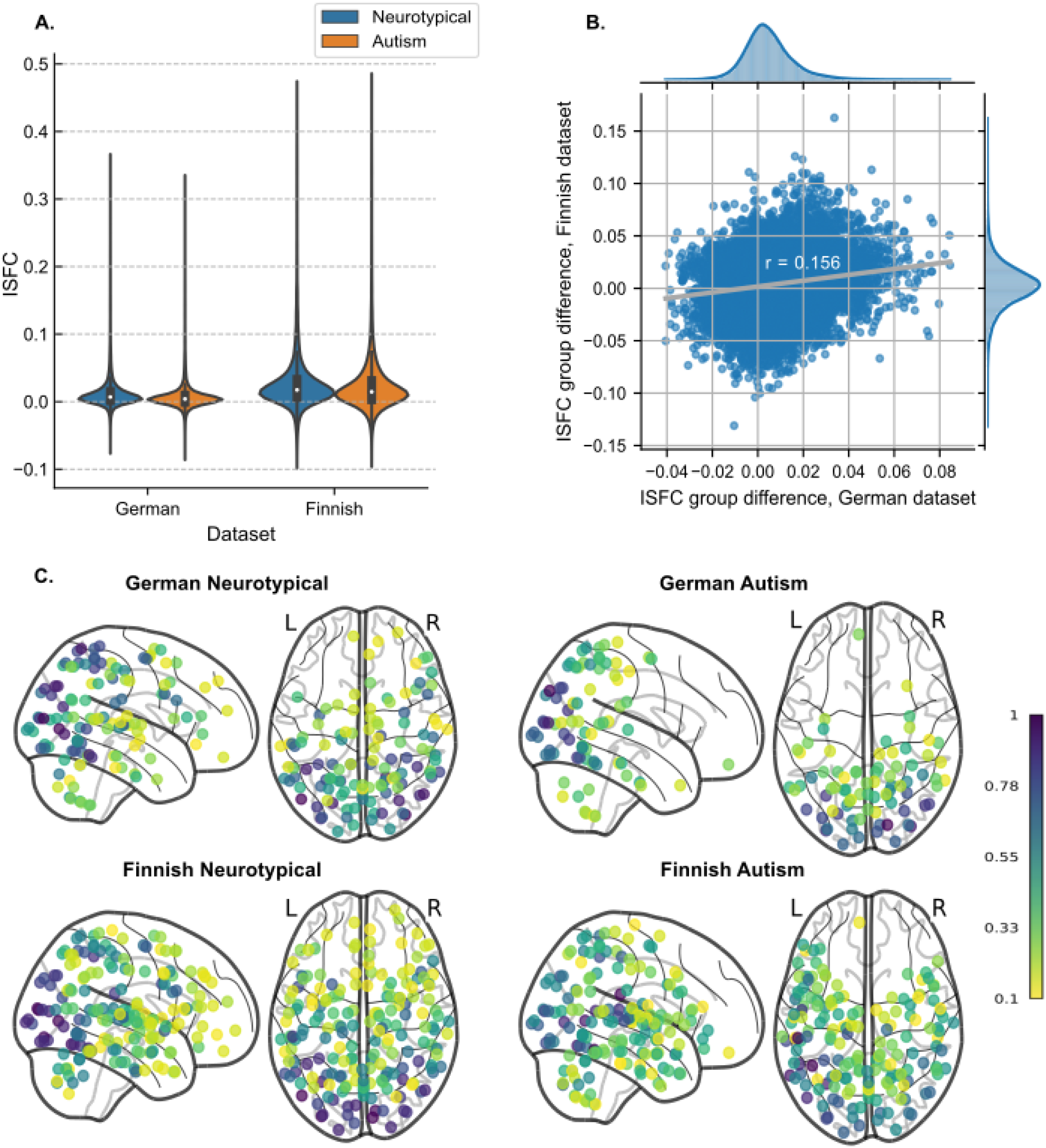
ISFC distribution. (A) Violin plots of pairwise median ISFC values over pairs of regions across each group (neurotypical, autism) and each dataset (German, Finnish). The whit dot in each box plot within the violin plot represents the group’s median ISFC value. (B) Scatterplot of ISFC difference (neurotypical-autism) in the German and Finnish dataset. ISFC values are pairwise median values across neurotypical and autism participants in each dataset. The distributions of the ISFC differences are shown at the top for the German dataset and on the right for the Finnish dataset. (C) Count maps of ROI regions with significant pairwise ISFC across four subgroups, p_permuted_ < .001. The color of each dot represents the normalized count as a ratio to the maximum count within its specific group. Only ROI regions with a normalized count of at least 0.1 were visualized.

Overall, the regions with lower ISFC in autism vs. neurotypical group were similar across German and Finnish datasets, focusing on temporo-occipital visual regions, parietal cortex, precentral and dorsomedial prefrontal cortex. By contrast, temporal regions showed the reverse ISFC group differences (autism>neurotypical) in the Finnish dataset, whereas no such differences were observed in the German dataset. To show the differences between groups across brain regions, we mapped the number of significant pairwise connections for each ROI onto bilateral brain surfaces (Figure 3A, B). Subcortical results are mapped in Figure S1. The detailed counts of all significant ROIs are listed in Supplementary Table 5.

**Figure 3.**
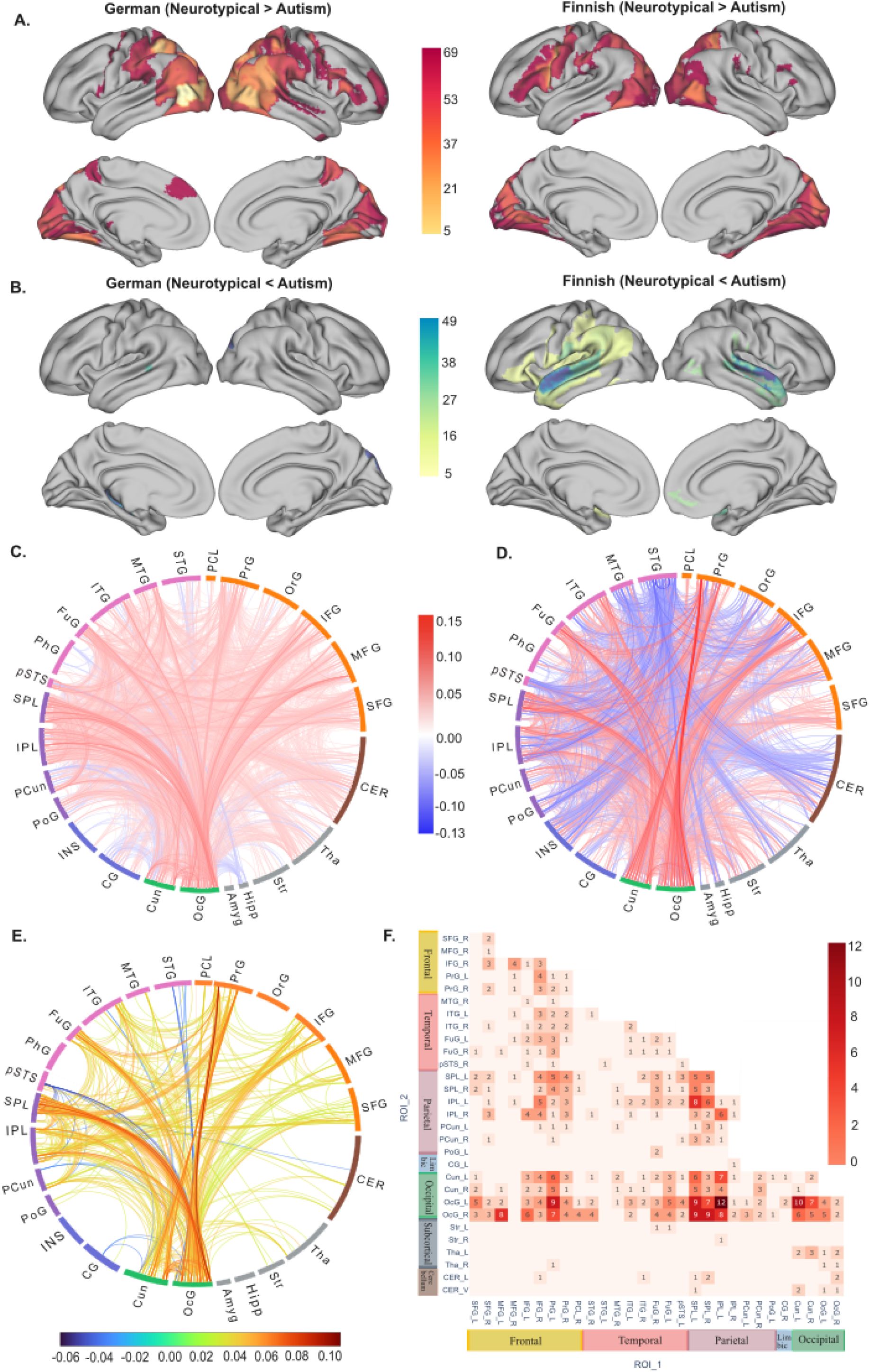
ISFC group difference. (A) Count maps of ROIs where neurotypical > autism, p < .01. The maximum positive count is 69 for the German dataset and 48 for the Finnish dataset. (B) Count maps of ROIs where neurotypical < autism, p < .01. The maximum negative count is 8 for the German dataset and 49 for the Finnish dataset. Only ROIs with at least 5 significant ISFC counts were visualized. (C) German ISFC group difference. (D) Finnish ISFC group difference.(E) The overlap of ISFC group differences, mapped with the average ISFC difference between neurotypical and autism groups across both datasets. The connectome maps were created using <u>NiChord</u> (41) (F) The counts of replicated ISFC group differences mapped in combined Brainnetome and Cerebellum atlas with a total of 273 ROIs, p < .05. Each annotated value indicates the number of significant pairwise ISFCs between the ROI regions listed on the x-axis and y-axis. SFG, Superior Frontal Gyrus; MFG, Middle Frontal Gyrus; IFG, Inferior Frontal Gyrus; PrG, Precentral Gyrus; PCL, Paracentral Lobule; STG, Superior Temporal Gyrus; MTG, Middle Temporal Gyrus; ITG, Inferior Temporal Gyrus; FuG, Fusiform Gyrus; pSTS, Posterior Superior Temporal Sulcus; SPL, Superior Parietal Lobule; IPL, Inferior Parietal Lobule; PCun, Precuneus; PoG, Postcentral Gyrus; CG, Cingulate Gyrus; Cun, Cuneus; OcG, Occipital Gyrus; Amyg, Amygdala; Hipp, Hippocampus; Str, Striatum; Tha, Thalamus; CER, Cerebellum. L, left hemisphere; R, right hemisphere; V, vermis.

ISFC group differences were similar between German males and Finnish males (Figure S2). No significant differences were found between ISFC group difference matrices before and after regressing out age and sex in the two datasets (Figure S3).

### 3.2 Replicate ISFC differences across datasets

We observed 3500 significant pairwise ISFC differences between the groups in the German dataset and 3058 pairwise differences in the Finnish dataset (two-sample subject-wise permutation test, two-tailed p < .05, uncorrected). 16% of the pairwise effects observed in the German dataset were replicated in the Finnish dataset (p_permuted_ < .001). The pairwise links in each dataset and the replicated connections are visualized in Figure 3C-E.

As shown in Figure 3F, the largest number of ISFC pairwise group differences were observed between the left Occipital Gyrus (OcG) and the left Inferior Parietal Lobule (IPL). The Superior Parietal Lobule (SPL) also correlated with ROIs in the visual regions including OcG and Cuneus. Besides, the Inferior Frontal Gyrus (IFG) and precentral gyrus showed many correlations with visual and parietal regions. Subcortically, ISFC differences were observed between Thalamus (Tha), Cerebellum and visual regions. Supplementary Table 6 contains the full list of overlapping pairwise ISFC differences across two datasets.

## 4 Discussion

In this cross-national fMRI study, we observed replicable reductions in inter-subject functional connectivity in autistic individuals while they viewed dynamic scenes with social and non-social content. Compared to the neurotypical controls, autistic individuals showed replicable inter-subject hypoconnectivity between the visual network and dorsal attention network in the occipital and parietal cortices during both visual-only and audiovisual conditions. Furthermore, inter-subject hyperconnectivity in the superior temporal regions was observed in autism only for audiovisual condition. These findings align with prior research on reduced neural similarity in autism (17-18, 23). Notably, the current results extend previous work by demonstrating replicable, stimulus-driven connectivity differences across multiple fMRI datasets and between visual and audiovisual modalities.

### 4.1 Inter-subject hypoconnectivity of occipitoparietal/frontal regions in autism

Compared to neurotypical controls, autistic individuals showed reduced inter-brain functional connectivity between occipital and parietal/frontal cortices during visual-only and audiovisual stimulation. The most robust replicable differences were observed between the visual network (e.g., Cuneus, OcG) and the attention network (e.g., IPL, SPL), as defined by the Yeo network (42).

Regions in the IPL, which belong to the dorsal and ventral attention networks, showed reduced ISFC in autistic individuals. These regions are critical for action observation and understanding the intentions of others (43,44). Decreased activity in the left IPL has been linked to impairments in attributing intentions to actions (45). Additionally, atypical IPL connectivity has been associated with deficits in social-communicative skills in autism (46). A previous study on prepubertal boys with autism reported reduced left IPL activity starting at an early age, potentially contributing to their social difficulties (47). Therefore, the reduced ISFC between the IPL and visual regions observed in this study may reflect impaired action understanding in autism, which could affect social cognition over time.

The autism group also exhibited reduced ISFCs between the bilateral SPL and visual regions. Large area of the SPL that align with the dorsal attention network were involved. This area plays a role in visual attention (48), action observation, and visuomotor integration (49). Prior research has shown decreased SPL activation during motor learning in autistic versus neurotypical individuals, with this reduction linked to repetitive behaviors (50). One potential explanation for the current findings is that autistic indviduals display idiosyncratic visual attention during movie-watching and struggle to integrate socially relevant sensorimotor cues (e.g., faces, body movements) compared to neurotypical individuals (17,51). After prolonged viewing, their visual attention to social stimuli was found to decline with no subsequent recovery, unlike neurotypical individuals (52).

In the frontal cortex, inter-subject hypoconnectivity was observed between the IFG and precentral gyrus with occipitoparietal regions. The IFG is involved in language processing (53), attention reorientation (54,55), inhibitory control (56,57), and social cognition (58). The precentral gyrus is essential for motor planning and execution (59). Reduced ISFC between the right IFG and occipitoparietal regions in autism may indicate less-regulated attention to visual stimuli in movie clips. Along with the SPL, decreased ISFC between the precentral gyrus and visual regions suggests altered coordination between visual input and motor output in autism. As parts of the putative mirror neuron system, the precentral gyrus, IFG and IPL participate in observing and imitating actions (60,61). Reduced ISFC between these regions and visual areas may contribute to the challenages autistic individuals experience in understanding and mimicking others’ actions, particularly during tasks that require social attention, such as movie-watching.

In temporal regions, the fusiform gyrus (FFG) exhibited widespread inter-subject hypoconnectivity with occipitoparietal regions. The FFG is known for face recognition (e.g., the fusiform face area, 62) and object color recognition (63). Its role in social functioning among autistic individuals has been extensively studied (64-66). As a key hub in social processing, reduced ISFC between the FFG and occipitoparietal regions observed here may underlie the social cognition impairments seen in autism.

In subcortical regions, our results demonstrated lower ISFCs between the thalamus/cerebellum and visual regions in autistic individuals compared to the neurotypical group. The thalamus is crucial for modulating sensorimotor signals and integrating sensory information with motor outputs (67). Dysconnectivity between the thalamus and various cortical regions, including prefrontal, temporal, and sensorimotor cortices, has been reported in autism (68,69). Similarly, the cerebellum is well-established in motor control (70,71) and sensorimotor functions (72,73). Recent research has expanded its role to social cognition, such as processing social interactions (74,75). The cerebellum may mediate links between sensorimotor processes and higher-level social-cognitive functions, although this role appears weaker in autistic individuals compared to neurotypical controls (75). However, the ISFC differences in the thalamus and cerebellum were smaller than those observed in cortical regions (e.g., parietal and frontal areas) connected to visual regions. This finding may suggest that subcortical regions exhibit less inter-subject variability, making it harder to detect group-level differences.

### 4.2 Inter-subject hyperconnectivity of temporal regions in autism

In the audiovisual condition, autistic individuals showed stronger ISFCs between temporal (e.g., pSTS, STG) and visual regions compared to neurotypical controls. This effect was not observed in the visual-only condition. Inter-subject hyperconnectivity in the left pSTS contrasts with prior findings of within-subject underconnectivity in parts of the pSTS in autism, which are more commonly reported for the right hemisphere (76,77). The pSTS plays a key role in receiving polymodal input and integrating convergent sensory processes (78), which may explain the aberrant inter-subject connectivity between temporal and visual regions in autism. In the initial study using the current stimuli, the bilateral pSTS showed widespread activation in response to the videos’ social content (32). Additionally, the social context of the movie stimuli can be classified from brain activity patterns in the social perceptual network, including the STS and STG (79). These findings suggest that autistic individuals may require more neural resources than neurotypical individuals to integrate sensory information in social contexts, potentially explaining the observed hyperconnectivity in autism.

In the visual-only condition, ISFCs from superior temporal regions were weaker. The absence of a soundtrack during the moving-watching task in the German dataset may have reduced ISFC in the temporal lobes and could also globally influence ISFC magnitudes. Maximal ISFCs were lower in the German dataset (visual-only) compared to the Finnish dataset (audiovisual), which may increase overall ISFC in three ways. First, local ISFCs connected to early auditory regions may increase due to the tight coupling of auditory cortical activity with acoustic features of the stimulus (e.g., 80). Second, temporofrontal language regions may follow the occurrence of speech during naturalistic stimulation (32,80). Third, in a naturalistic fMRI paradigm, the soundtrack likely enhances the movie-watching experience by facilitating attention synchrony(81) and multisensory integration (82). This enhanced engagement may elevate ISFC levels across different brain regions in audiovisual conditions.

### 4.3 Sex and cross-cultural asymmetry in autism

We observed smaller ISFC differences in autistic females compared to both Finnish and German males (see Supplemental results). The between-sex differences were comparable to between-country differences, suggesting that sex distribution had minimal impact on ISFC variation across countries. Sex differences have received increasing attention in autism research (83-85). Autism is more commonly diagnosed in males, with an estimated male-to-female ratio of 4:1 (84). Females are found to camouflage their autistic traits better, potentially leading to underdiagnosis (87,88). Additionally, autism in females is often associated with fewer social impairments, such as communication difficulties, compared to males (89). The varied diagnosis criteria across cultural contexts can introduce additional variability when studying inter-subject neural similarity. Culture shapes individuals’ social communication and cognition (90), potentially affecting inter-subject variability in neural responses across different populations (91,92).

### 4.4 Limitations

The current replication results should be interpreted with its limitations. First, the current study used subject-wise permutation (SWP) without additional statistical control for multiple comparisons. Chen et al. (40) showed SWP to be the most effective approach for two-sample tests in whole-brain voxel-wise analyses of ISC, outperforming other non-parametric methods like element-wise permutation and subject-wise bootstrapping in controlling false positive rates (see their Figure 2). The BrainIAK toolbox used in the current study was rigorously developed based on Chen’s work, giving us confidence in its ability to limit false positives. However, future validation with simulated and real datasets, using larger sample sizes and ROI configurations, is needed to assess SWP’s robustness.

Additionally, replication challenges in autism research remain significant due to the wide variability in clinical symptoms across individuals (4) and cultures (90,91), which likely reflect their diverse neural correlates such as ISFC. Previous research has shown a lack of replication in functional connectivity differences between autism and neurotypical groups across multi-site datasets (93). Factors such as differences in imaging sites, methodological approaches (e.g., voxel-wise vs. region-specific analyses), and data analysis flexibility may contribute to the replication challenges (25). Furthermore, large sample sizes are critical for replicable results in task-based fMRI studies (94). Our datasets were collected as parts of two separate studies limiting the size of data and leading to differences in sampling and behavioral measures. More systematic collaboration across multiple sites using the same naturalistic paradigm holds promise for a more robust evaluation of the replication of group differences between autistic and neurotypical individuals.

Finally, the inter-subject analyses in this study focused on stimulus-driven brain activity, differing from assessments of intrinsic functional connectivity replicability. Future research should evaluate the replicability of both extrinsic and intrinsic contributors to group differences in functional connectivity in larger cohorts.

### 4.5 Conclusion

Our study highlights the idiosyncrasies of brain activity in autistic individuals compared to neurotypical group by examining their inter-subject functional connectivity during naturalistic movie-watching. The results demonstrated replicable inter-subject hypoconnectivity between visual, posterior temporal, and parietal regions in autism across two countries and two stimulation conditions. Similar effects were also observed when male and female participants were analyzed separately. Additionally, inter-subject hyperconnectivity in superior temporal regions was observed during the audiovisual condition in autism. These findings underscore atypical sensory and attentional processing of naturalistic stimuli in autism, emphasizing the potential of ISFC and naturalistic fMRI for detecting stimulus-driven neural connectivity changes associated with neurological and psychiatric conditions.

## Supporting information

Supplemental results

## Acknowledgments and Disclosures

This work was supported by the Finnish Cultural Foundation (Grant No. 150496 [to JML]), and the Independent Max-Planck-Research Group by the Max-Planck-Society (to LS). The authors report no biomedical financial interests or potential conflicts of interest.

