## Supplemental results for "Reduced inter-subject functional connectivity during movies in autism: Replicability across cross-national fMRI datasets"

**Description of clinical scores:**

In the German data, all participants completed the Autism-spectrum Quotient (AQ) questionnaire (1), the Anticipatory and Consummatory Interpersonal Pleasure Scale (ACIPS) (2,3), the Liebowitz Social Anxiety Scale (LSAS) (4), the Toronto Alexithymia Scale (TAS-20) (5), the Beck’s Depression Inventory (BDI-II, German version) (6), the Reading the Mind in the Eyes test (RME) (7). Most German participants were assessed using the Autism Diagnostic Observation Schedule-2 (ADOS-2) – Module 4 (8) by experienced clinical psychologists. In the Finnish data, all participants completed the AQ questionnaire, the BDI-II (9), the Depression Anxiety Stress Scale-21 (DASS-21) (10), and the State-Trait Anxiety Inventory-X2 (STAI-X2) (11). ADOS-2 assessment was also performed by a trained clinical psychologist to quantify the autism severity in the autism group (12). All group scores described above are listed in Tables S1-4.

The ADOS assessment used in the German and Finnish datasets was based on the same version designed for adult autistic individuals, but administered in different languages. ADOS scores in both autism groups were significantly different from the within-site neurotypical group. However, a Mann-Whitney U test indicated that ADOS scores in the German autism group were significantly lower than those in the Finnish autism group (U = 63.5, *p* = 0.0018, *r* = -0.519). This difference may reflect variations in diagnostic criteria and procedures across countries. Conversely, AQ scores were significantly higher in the German autism group compared to the Finnish autism group (U = 342.0, *p* = 0.0001, *r* = 0.619). Since the AQ is a self-report measure, these results may be influenced by subjective bias. Cultural differences in self-perception may also play a role. However, these potential cross-cultural differences cannot be directly quantified within the current datasets.

**Table S1.** Clinical scores of the German Autism group.

|  | **ADOS** | **AQ** | **ACIPS** | **LSAS** | **TAS-20** | **BDI_II** | **RME** |
| --- | --- | --- | --- | --- | --- | --- | --- |
| **N** | 18 | 22 | 22 | 22 | 22 | 22 | 22 |
| **Mean** | 7 | 37.59 | 52.09 | 74.25 | 59.93 | 12.45 | 19.68 |
| **Std** | 3.83 | 7.96 | 13.20 | 21.57 | 7.88 | 9.71 | 4.81 |

**Table S2.** Clinical scores of the German Neurotypical group.

|  | **ADOS** | **AQ** | **ACIPS** | **LSAS** | **TAS-20** | **BDI_II** | **RME** |
| --- | --- | --- | --- | --- | --- | --- | --- |
| **N** | 14 | 25 | 25 | 25 | 25 | 25 | 25 |
| **Mean** | 0.78 | 13.68 | 84 | 20.96 | 40.68 | 4.12 | 25.44 |
| **Std** | 1.05 | 5.61 | 8.50 | 14.27 | 9.98 | 5.14 | 3.27 |

**Table S3.** Clinical scores of the Finnish Autism group.

|  | **ADOS** | **AQ** | **BDI-II** | **DASS-21** | **STAI-X2** |
| --- | --- | --- | --- | --- | --- |
| **N** | 18 | 18 | 18 | 18 | 18 |
| **Mean** | 12 | 27.44 | 8.22 | 14.88 | 43.83 |
| **Std** | 4.32 | 5.86 | 7.36 | 8.93 | 8.11 |

**Table S4.** Clinical scores of the Finnish Neurotypical group.

|  | **AQ** | **BDI-II** | **DASS-21** | **STAI-X2** |
| --- | --- | --- | --- | --- |
| **N** | 19 | 19 | 19 | 19 |
| **Mean** | 10.94 | 3.05 | 7.52 | 36.47 |
| **Std** | 3.53 | 2.91 | 5.96 | 6.33 |

**Comorbidities and medicine of autistic groups:**

**Table S5.** Diagnosis and medical status of the German Autism group.

| **Subject** | **Diagnosis** | **Medication** |
| --- | --- | --- |
| 1 | ASD | None |
| 2 | ASD | None |
| 3 | ASD, MDD | Antidepressants |
| 4 | ASD, MDD, SAD | Antipsychotics |
| 5 | ASD, MDD | Antidepressants,  Antipsychotics |
| 6 | ASD | None |
| 7 | ASD, MDD, OCD | Antidepressants,  Antipsychotics |
| 8 | ASD, MDD | None |
| 9 | ASD | None |
| 10 | ASD | Antidepressants |
| 11 | ASD | None |
| 12 | ASD | Lorazepam |
| 13 | ASD, MDD | Antidepressants |
| 14 | ASD | None |
| 15 | ASD, MDD | Antidepressants |
| 16 | ASD | Antipsychotics |
| 17 | ASD, MDD | Antidepressants |
| 18 | ASD | None |
| 19 | ASD | None |
| 20 | ASD, MDD, SAD,  Dyslexia | Antidepressants,  Antipsychotics |
| 21 | ASD | None |
| 22 | ASD, SAD | None |

*Note: MDD = Major Depressive Disorder.*

**Table S6.** Diagnosis and medical status of the Finnish Autism group.

| **Subject** | **Diagnosis** | **Medication** |
| --- | --- | --- |
| 1 | ASD, MAD | None |
| 2 | ASD | None |
| 3 | ASD, MAD | None |
| 4 | ASD, ADHD | Melatonin |
| 5 | ASD, MAD | Fluoxetine |
| 6 | ASD | None |
| 7 | ASD, ADHD | Zolpidem |
| 8 | ASD, ADHD | None |
| 9 | ASD, MAD | Melatonin |
| 10 | ASD | None |
| 11 | ASD, ADHD | None |
| 12 | ASD, MAD | Venlafaxine |
| 13 | ASD | None |
| 14 | ASD | None |
| 15 | ASD, MAD | Escitalopram |
| 16 | ASD, MAD | Vortioxetine, Bupropion (stopped 5 days before) |
| 17 | ASD, ADHD | Melatonin |
| 18 | ASD, ADHD | None |

*Note: MAD = Mood and Anxiety Disorder (excluding*

*severe mental disorders by SCID-I).*

**Figure S1.** Count of subcortical regions with ISFC group difference (two directions, p < .01). Amyg: Amygdala; Hipp: Hippocampus; Str: Striatum; Tha: Thalamus; CER: Cerebellum. Details with region names and values are listed in Supplementary Table 5.

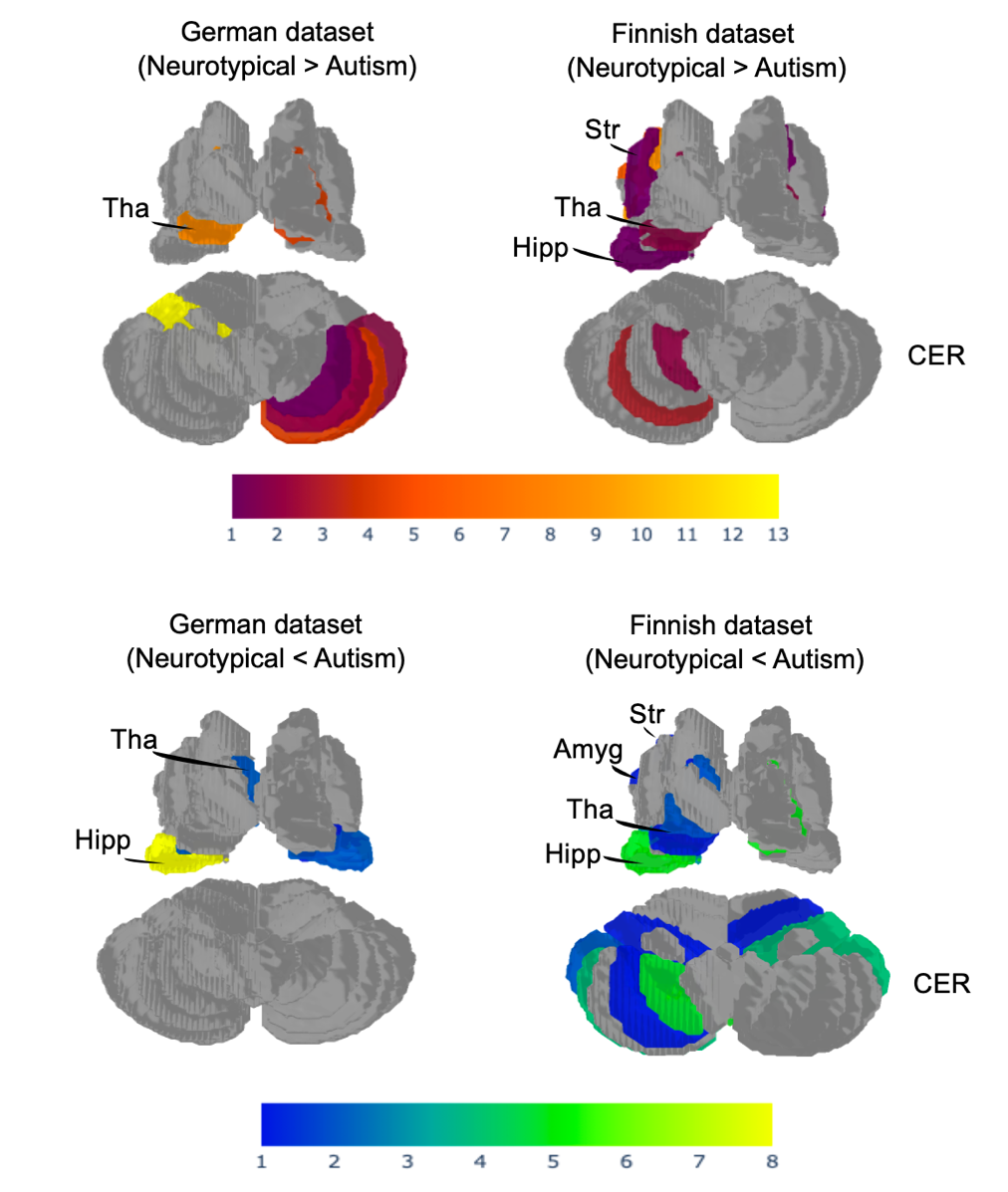

**ISFC group differences across sexes in two datasets**

We observed higher correlation and replication rates of pairwise ISFC group differences across sexes than across countries (Figure S2). The replication rate between German males and females was 19% at an ISFC significance level of p < .05. ISFC differences showed significant correlations between German males and females (Pearson r = 0.201, p_permuted_ < .001) and between German males and Finnish males (Pearson r = 0.146, p_permuted_ < .001). However, German females showed a weak correlation with Finnish males (Pearson r = 0.022, p_permuted_ = .130). Overall, males exhibited higher ISFCs than females in both diagnostic groups. Notably, ISFC differences between neurotypical and autistic individuals in Finnish males more closely resembled those of German males than those of German females.

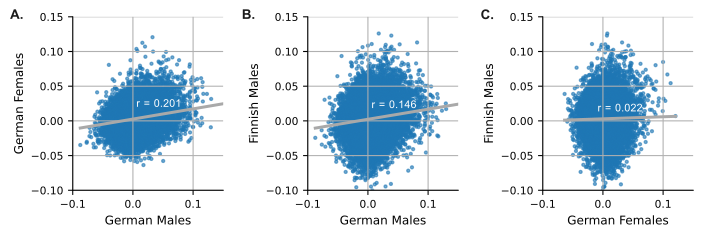

**Figure S2**. Scatterplots of pairwise ISFC difference (neurotypical-autism) across sexes in German and Finnish participants. (A) German males vs. German females. (B) German males vs. Finnish males. (C) German females vs. Finnish males. The ROI orders were randomly permuted 5000 times to generate the null distribution for the Pearson correlations between different groups.

**Effects of age and sex on the ISFC group differences**

The age distribution in the German and Finnish datasets shows slight differences, with the German dataset having a higher mean and standard deviation. Independent t-tests revealed no significant age differences between neurotypical and autism groups within each dataset, nor between the neurotypical groups across datasets. However, the German autism group was significantly older than the Finnish autism group (t = 3.360, *p* = 0.0019).

To evaluate potential confounding effects of age and sex on ISFC group difference, linear regression was used to remove these effects (where available) from the ISFC matrices. In the Finnish dataset, available confounders included pairwise mean age and absolute age difference. In the German dataset, sex was included as an additional confounder. All confound variables were z-scored, and a constant term was added. For each ROI pair, linear models were fit across subject pairs, and confound-related variance was regressed out, resulting in cleaned ISFC matrices that are orthogonal to age and sex while preserving the effects of different diagnostic groups.

Next, group-level median ISFC matrices were computed for neurotypical and autism groups, both before and after confound regression. No notable changes were observed. This was confirmed by a Mantel test comparing pre- and post-regression difference matrices, which showed a strong correlation (r = 0.98, *p* < 0.001, 5000 permutations), indicating that diagnostic effects were not primarily driven by age- or sex-related confounds (Figure S3).

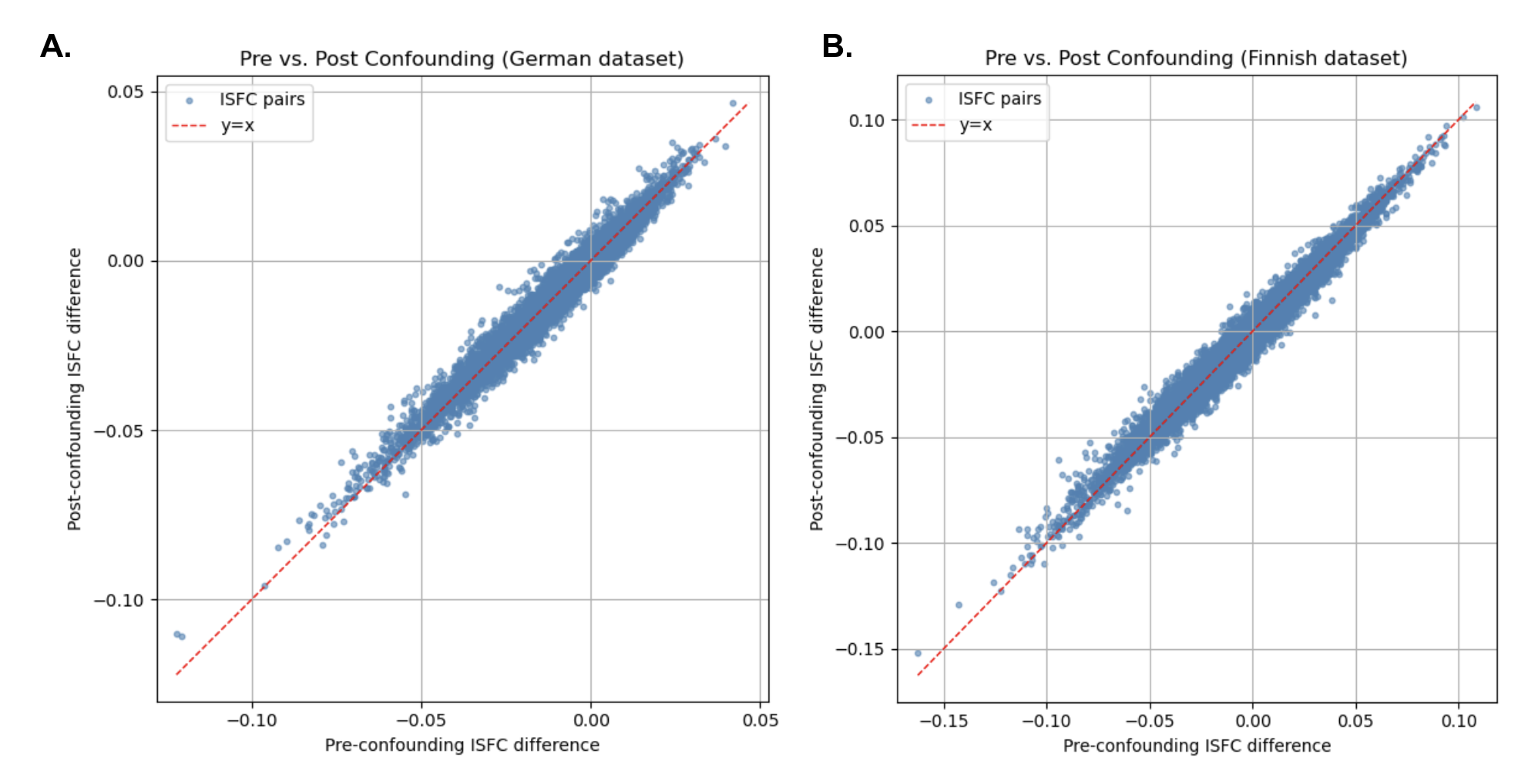

**Figure S3**. Scatterplots comparing group differences in ISFC before and after confound regression for the (A) German and (B) Finnish datasets. Each datapoint represents a median pairwise ISFC group difference (Neurotypical - Autism). The x-axis shows the uncorrected group difference, while the y-axis reflects the group difference after regressing out relevant confounds. A diagonal line is plotted in red for comparison of pre- and post-regression values.
